# Identification of GEFs and GAPS modulating phosphorylation and abundance of Rab10 by *LRRK2-G2019S* in neurons

**DOI:** 10.1101/2023.05.23.541929

**Authors:** Alison Fellgett, Sean T. Sweeney, Sangeeta Chawla, Christopher J. H. Elliott

**Author notes:** Correspondence to Christopher J. H. Elliott.

## Abstract

The Parkinson ‘s Disease associated kinase LRRK2 is proposed to act through phosphorylation of Rab proteins, most notably Rab10. All Rabs participate in a GTPase cycle, in which GEFs (Guanine nucleotide exchange factors) promote membrane binding, and GAPs (GTPase-activating proteins) promote release into the cytoplasm. LRRK2 is proposed to phosphorylate membrane bound Rab10. The hypothesis is that phosphoRab10 is less sensitive to GAP action and may remain membrane bound. To test how LRRK2 and Rab10 function in the GTPase cycle, we used an immunoblotting strategy in fly brains to show that a putative GEF Crag/DENND4C and three possible GAPs (pollux (plx), GAPcenA and Evi5, orthologs of AS160) interact with LRRK2 controlling the phosphorylation and abundance of Rab10. Crag behaves similarly to a Rab10 GEF and additionally modulates the level of panRab10. Only plx acts as a conventional GAP. GAPcenA seems to act as a GAP for phosphoRab10 more than panRab10. It is likely that Evi5 acts as a GAP for another Rab, possibly Rab11.

## Introduction

A common cause of Parkinson ‘s is the *G2019S* mutation in *LRRK2*. This amino-acid change increases the kinase activity of LRRK2 with consequent hyperphosphorylation of several GTPases. One of the best studied of these GTPase is Rab10, which shows hyperphosphorylation under the expression of *LRRK2*-*G2019S* in cell culture (Ito et al., 2016). Like other GTPases, Rab10 cycles between a membrane bound, active GTP-Rab10 and cytosolic, inactive, GDP-bound form (Fig. 1). Phosphorylation may affect the binding of effectors, cellular protein locations and rate of this cycle. Importantly, it has been suggested that phosphorylated Rabs cannot be released from the membrane to the cytosol (Liu et al., 2018). This indicates the importance of determining which GEFs (guanine nucleotide exchange factors) and GAPs (GTPase-activating proteins) operate to modulate the Rab10-GTPase cycle, and to see how these are affected by *LRRK2*-*G2019S*. This is a key question, as GEFs and GAPs may bind with multiple Rabs and/or interact with a sequence of Rabs.

**Fig. 1.**
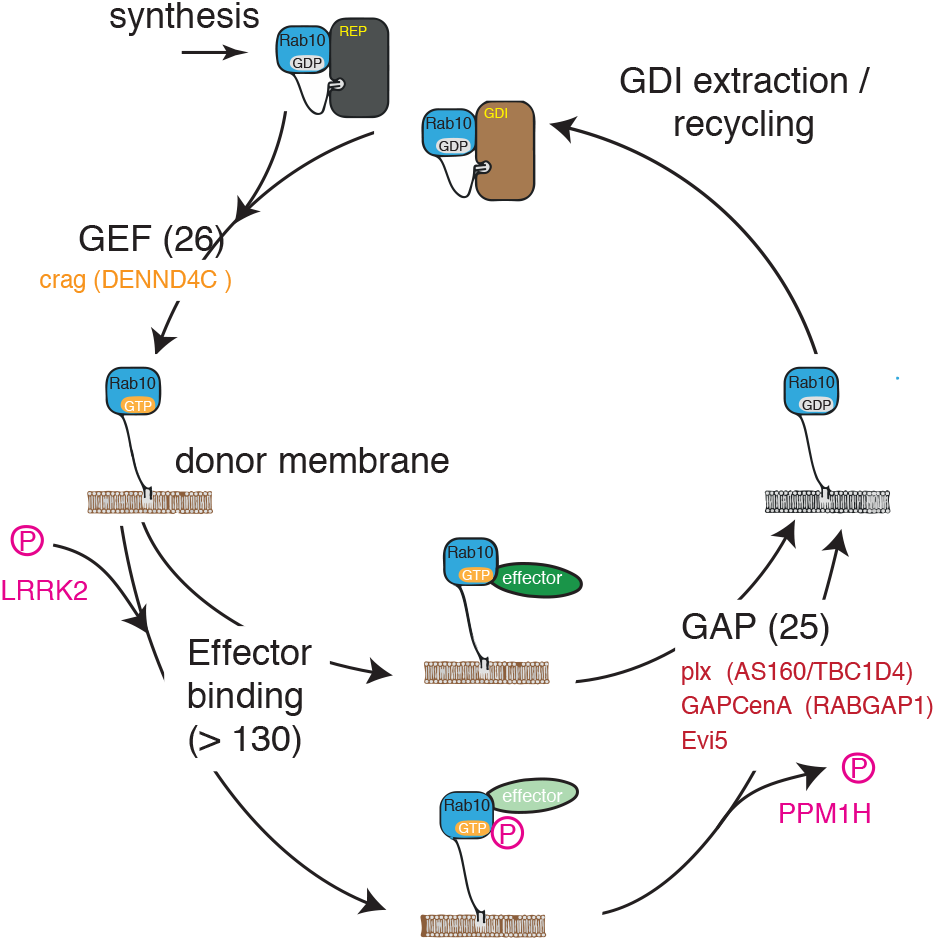
Proposed GEF– GAP cycle for Rab10 and LRRK2. After synthesis and prenylation by Rep, a GEF (guanine nucleotide exchange factor) facilitates the binding of GTP to Rab10. GTP-Rab10 is then inserted in a donor membrane. There are 26 potential GEFs in Flybase (http://flybase.org/reports/FBgg0000337.html), of which only one (Crag) is an ortholog of the hRab10 GEF DENND4C. Once inserted in the membrane, GTP-Rab10 may be phosphorylated by LRRK2, and this alters the effector(s) being bound. There are more than 130 potential effectors for Rabs (Gillingham et al., 2014). Under the action of a GAP (GTPase-activating protein), Rab10 reverts to a GDP bound form. In the fly, there are 25 GAPs (http://flybase.org/reports/FBgg0000331.html) of which 3 (plx, GAPcenA/RabGAP1, Evi5) may be orthologs of the hRab10 GAP TBC1D4/ AS160. Non-phosphorylated GDP-Rab10 leaves the membrane for the cytoplasm. This process is facilitated by the GDP dissociation inhibitor (GDI, a single protein in the fly). phosphoRab10 may remain membrane bound (Liu et al., 2018) until rescued by the PPM1H phosphatase (Berndsen et al., 2019). Proteins in brackets are the known interactors of hRab10, and orthologs of the fly proteins tested in this paper. Modified after (Goody et al., 2017; Taylor and Alessi, 2020).

In mammalian adipocytes, Rab10 has been well defined, regulating the translocation of GLUT4 (glucose transporter) to the plasma membrane. In this system the GEFs and GAPs for Rab10 have been identified as DENND4C (Sano et al., 2011) and AS160/TBCD4 (Sano et al., 2007) respectively. The fly dRab10 has a very similar sequence to hRab10 and is also phosphorylated by *hLRRK2*-*G2019S* expression in the fly brain (Petridi et al., 2020). More specifically, manipulations of *LRRK2*-*G2019S* and *Rab10* in fly dopaminergic neurons indicate a key role for phosphoRab10 in vision and in a simple reflex reaching movement (proboscis extension) (Fellgett et al., 2021).

Here we use fly genetics to show that a GEF (Crag, the ortholog of DENND4C) and three GAPs - pollux (plx), GAPcenA and Evi5, orthologs of AS160 - interact with LRRK2 controlling the phosphorylation of Rab10. Crag behaves like a Rab10 GEF and additionally modulates the level of panRab10, while plx alone acts as a conventional GAP.

## Methods

### Flies and Genetics

Expression was achieved in *Drosophila melanogaster* using the GAL4-UAS system for tissue specificity. The *nSyb*-GAL4 (gift of Julie Simpson, 2nd chromosome) was used to drive expression of UAS lines obtained from Bloomington (UAS-*Crag*, 58464; UAS-*Crag*-RNAi, 53261; UAS-*pollux*, 83635; UAS-pollux RNAi, 66313; UAS-*Evi5*-RNAi, 38350; UAS-*GAPcenA*-RNAi, 34976) or were from Gregory Emery, (UAS-*GAPcenA*, O/O59; UAS-*Evi5* wild-type, O/102; UAS-*Evi5R160A* mutant O/108), Wanli Smith (UAS-*LRRK2*-*G2019S*) or Cheng-Ting Chien (UAS-*LRRK2-G2019S-K1906M*). A wild-type control (+/ +) was made by crossing our laboratory stocks of *CS and w*_*1118*_. We also used a second control (*nSyb*/+) in which no transgene was expressed, by crossing *nSyb*-GAL4 with an empty vector line (Bloomington 36303). Routine fly genetics was used to generate crosses in which either no, one or two UAS genes were expressed. All crosses were viable, were raised at 25 °C, and heads collected and frozen on day 3 of adult life.

### Western Blot

were performed as described recently (Fellgett et al., 2021) using a mouse pan-Rab10 antibody (Nanotools, clone 605B11, 1:500, with Alexa 633 secondary, 1:500) and a rabbit phospho-Rab10 antibody (Abcam, ab230261, 1:1000, Alexa 488 secondary, 1:500). The polyclonal rat Crag antibody (Denef et al., 2008) was a kind gift from Olivier Devergne, Illinois University and used at 1:500. Our rabbit anti-synaptotagmin antibody was used as a loading control (West et al., 2015). The images collected on an Invitrogen iBright FL1000. Images were scanned and measured using ImageJ.

### Bioinformatics

Sequences were drawn from the NCBI Protein or UNIPROT databanks, and compared with NCBI Blast, using default settings. Conserved residues and trees were calculated in Jalview and using the tools at https://www.phylogeny.fr. For orthologs, a threshold of 30 % identity was required.

## Results

### LRRK2-G2019S increases and LRRK2-G2019S-K1906M decreases phosphoRab10

To generate an *in vivo* model where we could test the role of LRRK2 function on Rab10 phosphorylation status and abundance, we used the Drosophila GAL4/UAS system to express the activated form of LRRK2-G2019S in neurons using the pan-neuronal promoter *n-syb*-GAL4. Pan-Neuronal expression of *LRRK2*-*G2019S* with *nSyb-*GAL4 increased the level of phosphoRab10/panRab10 in the fly brain (Fig. 2,4,5,6,) in agreement with our previous observations (Fellgett et al., 2021). The separation of phosphoRab10 from panRab10 is clearly seen in the immunoblots, where the green band for phosphoRab10 migrates at an apparent higher molecular weight to the magenta band for panRab10. This difference in weight is characteristic of the extra charge from the phosphate group. We note that it was not necessary to use a ‘Phos-tag ‘ gel (Ito et al., 2016) in order to separate phosphoRab10 and panRab10. In all blots the ratio of phosphoRab10/panRab10 appears small as there is no detectable panRab10 signal under the phosphoRab10 band.

**Fig. 2.**
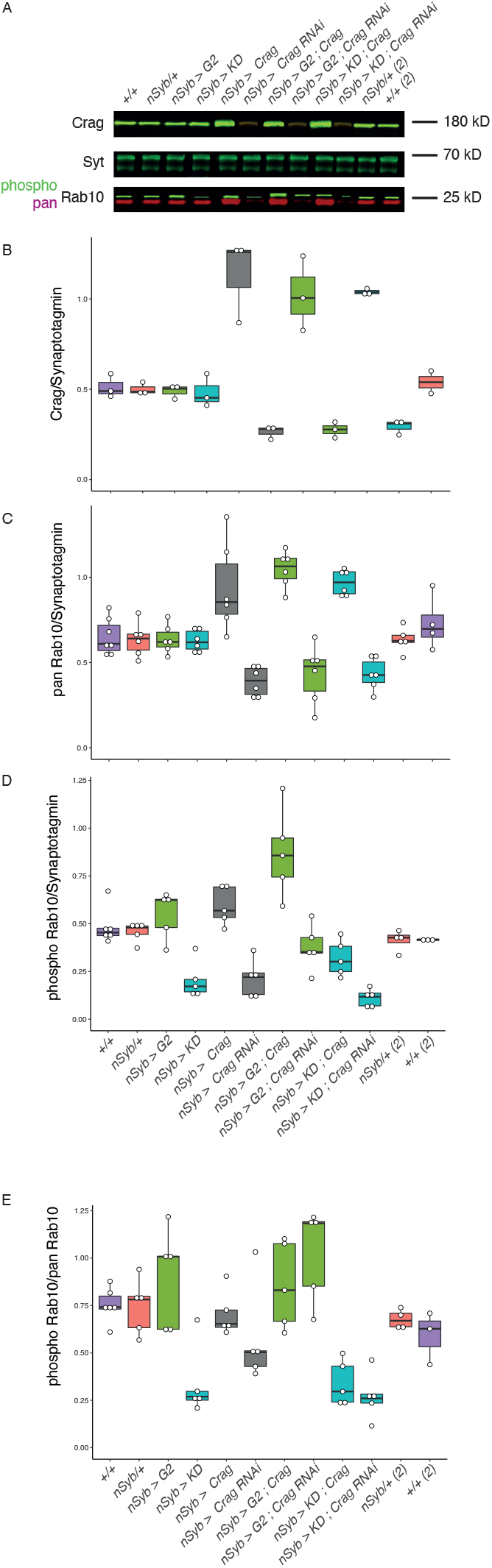
Crag acts as a GEF for Rab10 in drosophila CNS. A. Western blot showing the level of Crag, phosphoRab10 and panRab10, along with the loading control Synaptotagmin 91 (Syt). Manipulating LRRK2 had no effect on Crag, but the amount of Crag staining was increased by overexpression and reduced by *Crag-*RNAi. Increased Crag enhanced both phosphoRab10 and panRab10, with further increases in the *G2019S* background. Crag-RNAi or LRRK2-KD reduced phosphoRab10 with further reduction when both were expressed. These data suggest that Crag is acting as a GEF in both phosphoRab10 and non-phosphoRab10. Expression in the CNS was driven using *nSyb*. The wild-type control (+/ +) and no transgene-expressed-control (*nSyb/+*) were included at both ends of the blot. B. Quantification of the intensity of Crag/ Synaptotagmin in replicate blots. C. Quantification of the intensity of panRab10/Synaptotagmin in the replicate blots. D. Quantification of the intensity of phosphoRab10/Synaptotagmin in the replicate blots. E. Quantification of the intensity of phosphoRab10/ panRab10 in the replicate blots. Each dot represents one blot. Quantification of the Synaptotagmin loading control is shown in Suppl. Fig. 1. (*G2*: *LRRK2*-*G2019S*; *KD*: kinase dead LRRK2 (*LRRK2*-*G2019S-K1906M*).

Neuronal expression of the kinase dead (KD) form of LRRK2 (*G2019S-K1906M*) reduces phosphoRab10 abundance, suggesting it acts in a dominant negative manner.

### Expression of the GEF Crag/DENND4C increases both panRab10 and phosphoRab10

We hypothesise that increased expression of a Rab10 GEF should increase the amount of membrane-bound Rab10-GTP. In turn this allows for native kinases (e.g. dLRRK) or exogenous LRRK2 to phosphorylate Rab10. The fact that the proportion of Rab10 that is phosphorylated is quite small indicates that the phosphoRab10 provides a sensitive indicator of the effect of the GEF. Knock-down of the GEF should have the opposite effects.

There are 29 putative RabGEFs, of which 5 contain DENN domains in the fly genome. We find that only one GEF, Crag, encoded by *CG12737*, has an alignment with more than 45 % identity to the hRab10 GEF, *DENND4C*. The Crag-specific antibody detects a single band in fly heads at the expected Molecular Weight (187 kD), with the intensity of the band increased by neuronal overexpression of *Crag* and partially reduced by *Crag-*RNAi (Fig. 2A,B). CNS expression of the enhanced kinase form of LRRK2, *G2019S* or the kinase-dead form (*KD*) has no effect on the level of Crag (Fig. 2A,B).

Overexpression of Crag however increased the level of panRab10 above that of wild-type controls, while the RNAi reduced panRab10, irrespective of the level of LRRK2 kinase (Fig. 2C). We expect the panRab10 antibody to report both membrane-bound and cytoplasmic Rab10 as the antibody binding site (DDKRVV) is far from the prenylated tail, and from the switch II region where phosphorylation occurs. Thus the increased level of panRab10 when Crag is overexpressed, and concomitant decrease with Crag knock-down, suggests that Crag is a key component for the recruitment and abundance of Rab10.

Crag expression also increased phosphoRab10, while Crag-RNAi reduced it (Fig. 2D). To test for genetic interactions, we expressed both a *LRRK2* and a *Crag* transgene at the same time. When both Crag and G2019S levels were increased, the level of phosphoRab10 was greater than when just one of these proteins was manipulated. Conversely, when both Crag and KD proteins were reduced, the level of phosphoRab10 was less than when just one of these proteins was reduced Thus there appears to be a synergistic interaction between LRRK2 and Crag.

We also calculated the phosphoRab10/panRab10 ratio for each genetic manipulation. This ratio is increased with *G2019S*, and decreased by KD, but is not affected by Crag manipulations (Fig. 2E). This is the consequence of Crag affecting the levels of both phosphoRab10 and panRab10.

Our data is as expected for a Rab10 GEF but also indicates Crag may have a role in controlling the level of panRab10, possibly affecting the synthesis pathway.

### The GAPs pollux, GAPcenA and Evi5 interact differentially to regulate Rab10

Our hypothesis is that knock-down of a GAP will disrupt the GTPase cycle, by reducing the removal of Rab10 from the membrane, increasing the membrane-bound Rab10 and this will allow more Rab10 to be phosphorylated. The phosphoRab10 will remain in the membrane and provide an increased abundance. Conversely, overexpression of a Rab10 GAP should reduce the amount of phosphoRab10.

A BLAST search showed that, of the 25 GAPs in the fly, pollux (plx), GAPcenA and Evi5 were potential orthologs of the hRab10 GAP AS160/TBC1D4 (>30% identical residues, Fig. 3). Of these, plx had the greatest similarity to AS160.

**Fig. 3.**
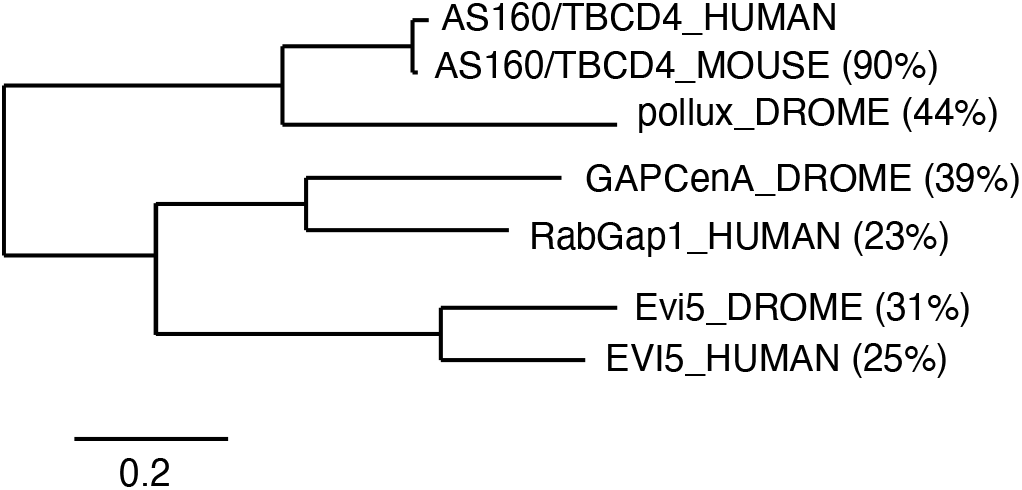
Phylogeny of GAPs linked to Rab10. Three *Drosophila* proteins, pollux, GAPcenA and Evi5 are potential orthologs of the hRab10 GAP AS160/TBCD4. pollux is the most similar, followed by GAPcenA and then Evi5. The human proteome also contains other orthologous GAPs (RabGap1, and EVI5). The percentages show the identity of each protein to AS160/TBCD4.

Pan-Neuronal expression of *plx* reduced the phosphorylation of Rab10 (Fig. 4A,C), while RNAi-mediated knock-down increased the phosphorylation of Rab10 above wild-type levels. This is as expected if plx is acting as a Rab10 GAP. We observed an additive effect when both *plx* and *LRRK2* were manipulated. Thus *G2019S* or *plx*-RNAi expression increased phosphoRab10, and expression of both together further increased phosphoRab10 abundance.

**Fig. 4.**
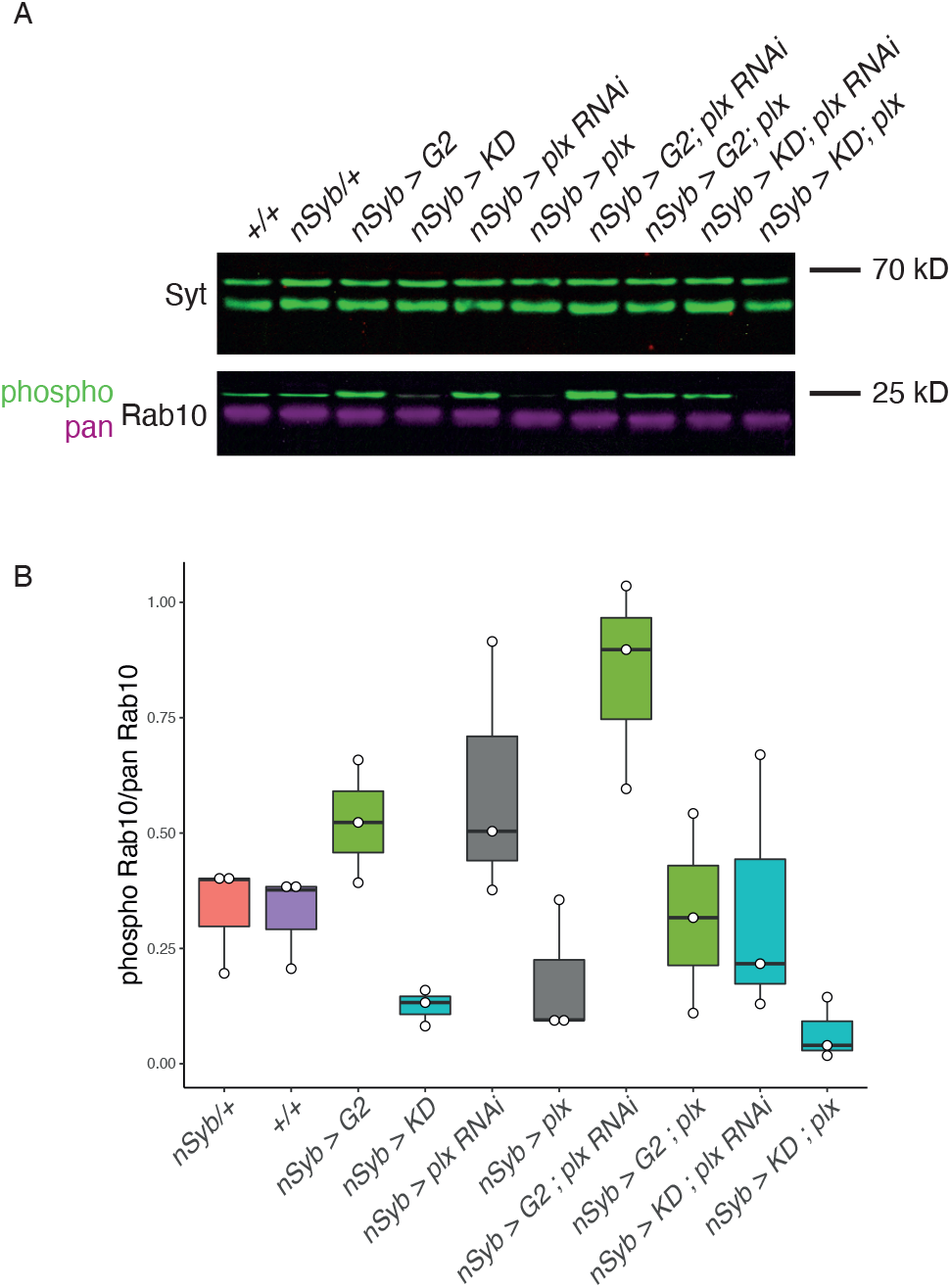
plx (pollux) acts as a GAP for Rab10 in drosophila CNS. A. Western blot showing the levels of phosphoRab10, panRab10 and the loading control Synaptotagmin 91 (Syt). *plx*-RNAi increased phosphoRab10 with further increase in the *G2019S* background. Increased *plx* expression reduced phosphoRab10, with further decrease in the *KD* background. These responses suggests that plx is acting as a GAP for both phosphoRab10 and non-phosphoRab10. There was no change in the panRab10/Synaptotagmin ratio (Suppl. Fig.2). Expression in the CNS was driven using *nSyb*. A wild-type control (+/+) and no transgene-expressed-control (*nSyb/+*) were included. B. Quantification of the intensity of phosphoRab10/ Synaptotagmin in the replicate blots. Data from 3 replicates. Quantification of the Synaptotagmin loading control is shown in Suppl. Fig. 1. (*G2*: *LRRK2*-*G2019S*; *KD*: kinase dead LRRK2 (*LRRK2*-*G2019S-K1906M*).

Similarly, either LRRK2-KD or plx reduced phosphoRab10, and both together further reduce phosphoRab10 abundance. Thus, this potential GAP effect applies to both phosphoRab10 and non-phosphoRab10. There was no effect of plx on panRab10 (Suppl. Fig. 2A).

The potential Rab10 GAP, GAPcenA, has a 39 % identity to AS160 (Fig. 3), and the human orthologs of GAPcenA include RabGap1 and RabGap1L. In the wild-type background, expression of *GAPcenA-*RNAi fails to produce the increase of phosphoRab10 expected if this protein is a Rab10 GAP (Fig. 5A,B). However, other features of the data are in line with the hypothesis that GAPcenA is a Rab10 GAP. These include the result of *GAPcenA*-RNAi expression in the LRRK2-*G2019S* background, which substantially increases phosphoRab10, potentiating the effect of *G2019S*. This might suggest that GAPcenA is acting as a GAP for phosphoRab10 rather than panRab10.

**Fig. 5.**
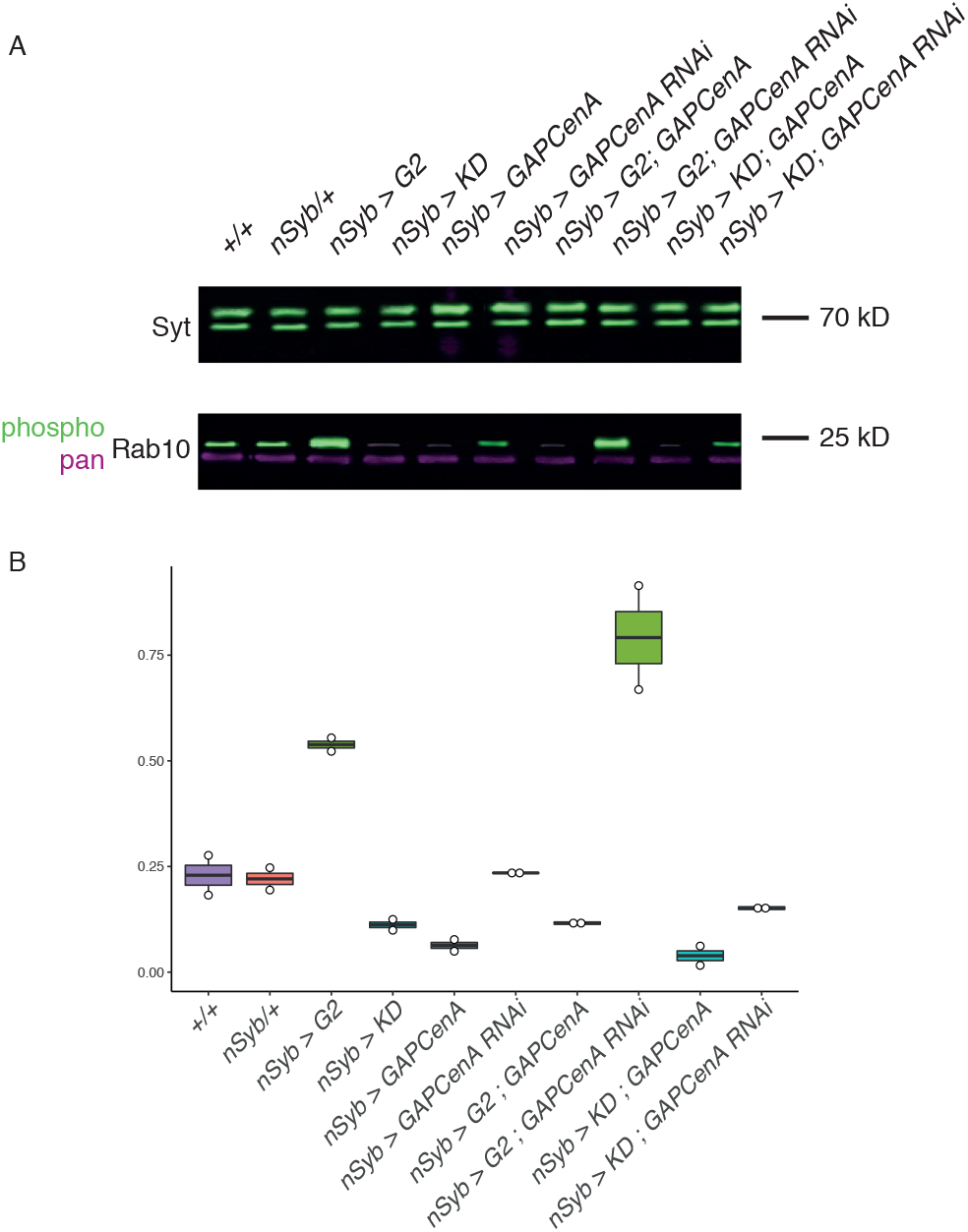
GAPcenA may act as a GAP for Rab10 in drosophila CNS. A. Western blot showing the levels of phosphoRab10, panRab10 and the loading control Synaptotagmin 91 (Syt). *GAPcenA*-RNAi resulted in little change from wild-type levels of phosphoRab10. With both *GAPcenA*-RNAi and *G2019S* a marked synergistic increase in phosphoRab10 was seen. Increased *GAPcenA* expression reduced phosphoRab10, with further decrease in the *KD* background. These responses suggest that GAPcenA may be acting more strongly as a GAP for phosphoRab10 than for non-phosphoRab10. There was no change in the panRab10/Synaptotagmin ratio (Suppl. Fig.2). Expression in the CNS was driven using *nSyb*. Controls included a wild-type (+/ +) and a no transgene-expressed-control (*nSyb/+*). B. Quantification of the intensity of phosphoRab10/Synaptotagmin in the replicate blots. Data from 2 replicates. Quantification of the Synaptotagmin loading control is shown in Suppl. Fig. 1. (*G2*: *LRRK2*-*G2019S*; *KD*: kinase dead LRRK2 (*LRRK2*-*G2019S-K1906M*).

Overexpression of *GAPcenA* reduced the amount of phosphoRab10 abundance in both the wild-type and *G2019S* backgrounds. In the KD background, the levels of phosphoRab10 are low, but suggest a small increase with GAPcenA RNAi and decrease with overexpression. There was no effect of GAPcenA on panRab10 (Suppl. Fig. 2B).

A third potential Rab10 GAP, Evi5, has a 31 % identity to AS160 (Fig. 3) and has two apparent orthologs in the human genome (EVI5, EVI5L). In the wild-type background, the phosphoRab10 levels were reduced by either neuronal expression of Evi5, Evi5 knock-down, or by expression of a catalytically dead form of Evi5 (*R160A*) (Fig. 6). In each case, the level of phosphoRab10 abundance was just above that seen in the KD brains. This is not the expectation from a Rab10 GAP. In the *G2019S* background, expression of wild-type or mutant *Evi5* did block the normal increase of phosphoRab10 - the level of phosphoRab10 was close to that of the wild-type. There was no effect of Evi5 on panRab10 (Suppl. Fig. 2C).

**Fig. 6.**
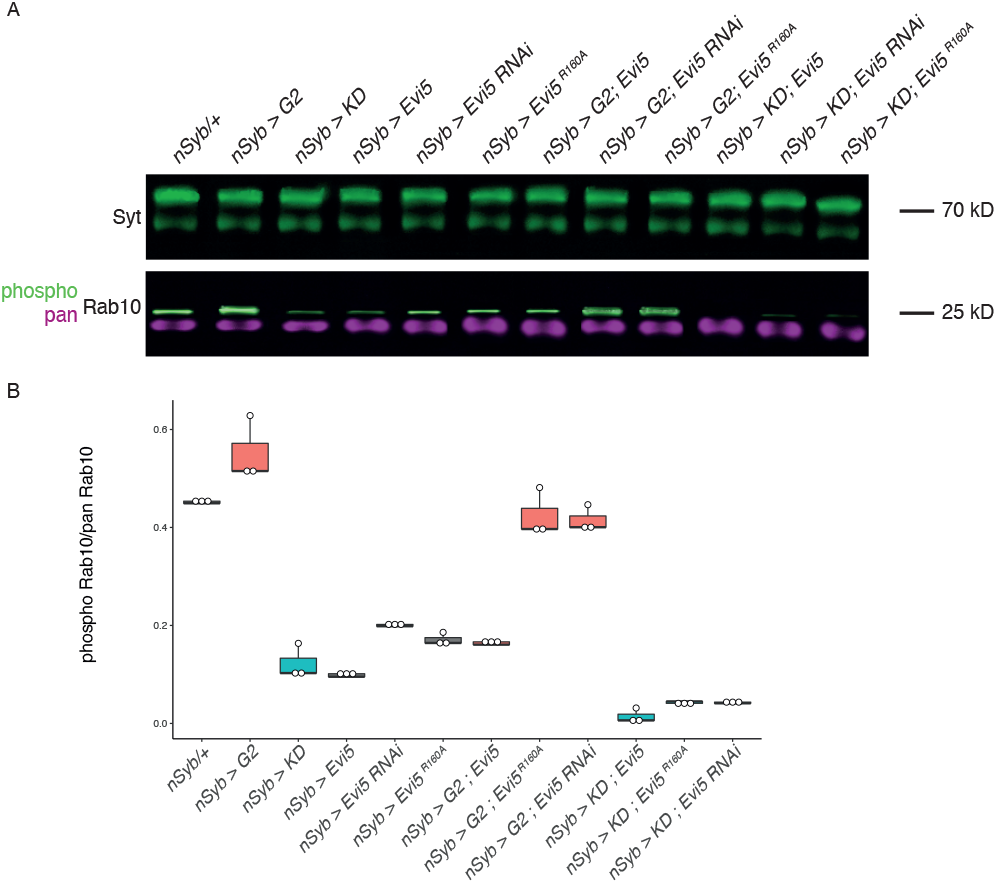
Evi5 does not act as a GAP for Rab10 in drosophila CNS. A. Western blot showing the levels of phosphoRab10, panRab10 and the loading control Synaptotagmin 91 (Syt). Expression of *Evi5*-RNAi, *Evi5* or catalytically inert *Evi5R160A* all reduced phosphoRab10. This is not the response expected of a Rab10 gap. There was no change in the panRab10/Synaptotagmin ratio (Suppl. Fig.2). Expression in the CNS was driven using *nSyb*. A wild-type control (+/+) and no transgene-expressed-control (*nSyb/+*) were included. B. Quantification of the intensity of phosphoRab10/ Synaptotagmin in the replicate blots. Data from 3 replicates. Quantification of the Synaptotagmin loading control is shown in Suppl. Fig. 1. (*G2*: *LRRK2*-*G2019S*; *KD*: kinase dead LRRK2 (*LRRK2*-*G2019S-K1906M*).

## Discussion

The Parkinson ‘s disease associated kinase LRRK2 is proposed to act through Rab10 action. We employed an in vivo system to test for GEFs and GAPs acting on Rab10 in the nervous system. Our main finding is that, in neurons, Crag and plx (pollux) potentially act as GEFs and GAPs for Rab10. Crag additionally acts on the recruitment of Rab10. While plx seems act as a GAP in all three backgrounds tested (wild-type, increased LRRK2 kinase, decreased kinase), GAPcenA acts as a GAP more strongly in the LRRK2-*G2019S* background (where there is more phosphoRab10) than in the wild-type. Evi5 does not seem to act as a Rab10 GAP.

### Crag as a GEF for Rab10

Our data demonstrates Crag meets the criteria of a GEF for Rab10 by promoting the levels of both panRab10 and phosphoRab10. In this, it facilitates the effect of LRRK2 on the Rab10 pathway.

The best understood role for Crag is in the secretion of basement membrane (BM), where it acts as a GEF for Rab10. Rab10 or Crag depletion results in secretion of the BM across both basal and apical surfaces of the Drosophila ovarian follicle cell (Lerner et al., 2013). In this system, loss of a different GEF, stratum (ortholog to RABIF), also affects secretion of the BM, but does this in a Rab8 dependent manner (Devergne et al., 2017). While the Crag/ Rab10 pathway is apically located, the stratum/Rab8 pathway is basally located.

Thus, in the follicle cell system, two independent pathways (which could both be LRRK2 sensitive), work to achieve the same effects. Crag-sensitive cell polarisation deficits have also been identified in fly photoreceptors. Here Crag, Rab10 and the effector Ehbp1 are required for the transport of the sodium/potassium ATPase from the Golgi to the basolateral membrane: a deficiency in one of the proteins results in translocation to the apical stalk membrane instead (Nakamura et al., 2020). Crag was also suggested to be a GEF for Rab11 (Xiong et al., 2012), but this idea depended in part on the belief that Rab10 was not expressed in fly neurons. More recent data indicate the presence of neuronal Rab10 (Davis et al., 2020; Fellgett et al., 2021; Kohrs et al., 2020), so we prefer a Rab10-Crag link. However, we also note that many Rabs work together in a sequential pathway, so a Rab11-Crag interaction is still possible.

### plx as a GAP for Rab10

We found that plx behaves exactly as expected of a GAP for Rab10. Increased plx expression reduces abundance of phosphoRab10 in neurons, and this potentially attenuates the effect of LRRK2.

Both Crag and plx are known to interact with calmodulin (Xu et al., 1998), but the function of this interaction is unclear. However, LRRK2 has been shown to affect calcium dynamics (Bedford et al., 2016), so this interaction may be important.

The human ortholog of plx, TBC1D4, exists as oligomers which increase the effectiveness of the interaction with Rab10 (Eickelschulte et al., 2021), and controls the translocation of GLUT4 to the plasma membrane as a response to insulin. Other Rabs (4A, 4B, 8A) are also associated with this pathway (Chen et al., 2012).

### GAPcenA as a partially effective GAP for Rab10

The role of GAPcenA is not fully established: we find it behaves as a Rab10 GAP for the phosphoRab10, rather than panRab10. This was unexpected as previous data suggests that phosphoRab10 remains membrane bound, and that GAPs are not very good at accelerating this process (Liu et al., 2018). In *Drosophila* larvae, GAPcenA attenuated the autophagy response (Zirin et al., 2015), which is intriguing as LRRK2 is frequently linked to autophagy.

Several Rabs have also been linked to autophagy, but (Zirin et al., 2015) did not identify a link between GAPcenA and Rab10.

### Evi5 may as a GAP downstream of Rab10

Our study suggests that Evi5 does not meet the criteria for a Rab10 GAP, as neither overexpression nor knock-down of Evi5 increase phosphoRab10.Nonetheless, our data indicates that Evi5 manipulations do affect the Rab10 GEF-GAP cycle. A possible explanation for this is that Evi5 is acting as a GAP for another Rab. It seems likely that this is Rab11, since Evi5 is the GAP for Rab11 in the fly ovary, where either overexpression or knock-down block cell migration (Laflamme et al., 2012). The human ortholog, EVI5 also interacts with Rab11a and Rab11b (Westlake et al., 2007). We suggest that disrupting the Evi5 GEF-GAP cycle prevents proper transfer of a Rab10 effector along a chain of membrane-bound Rabs.

### Linking GEFs or GAPs to LRRK2

Although many Rab10 effectors that might be downstream of LRRK2 have been identified, we are not aware of any previous evidence for GEF action in the LRRK2 - Rab10 pathway. Further, the results of LRRK2 -Rab10 GAP experiments in HEK293 cells are inconsistent (Fdez et al., 2022; Liu et al., 2018). The lack of data is surprising given the efforts to identify the main GEF and GAP for Rab10 in the insulin-signalling system. Only in the LRRK2 – Rab8 pathway has a potential GEF (Rabin8) and GAP (TBC1D15) been suggested (Steger et al., 2016). Here, using neuronal tissue, we confirm a role for the GEF Crag and GAP plx downstream of LRRK2, quite different from the LRRK2-Rab8 interactors.

Overall, the availability of specific antibodies for phosphoRab10 and panRab10 allow us to determine that Crag and plx disrupt the GTPase cycle as expected of a GEF and GAP, that the GAP GAPcenA seems to act on phosphoRab10 preferentially over panRab10.

Upstream of LRRK2, is another Rab, Rab29. This is atypical, with poor prenylation, interaction with membranes, and no GAP (Gomez et al., 2019). Nonetheless, in macrophage phagosomes, it does have a GEF (Rabaptin5), itself an effector of Rab5 (Shrivastava et al., 2022).

In conclusion, we note that the LRRK2 pathway has many GAPs and their roles generally require clarification.

## Figure legends

Suppl. Fig. 1. **Consistent levels of Synaptotagmin in all Western Blots**. Data from (A) Crag, (B) plx, (C) GAPcenA and (D) Evi5 blots. (*G2*: *LRRK2*-*G2019S*; *KD*: kinase dead LRRK2 (*LRRK2*-*G2019S-K1906M*).

Suppl. Fig. 2. **Consistent levels of panRab10/Synaptotagmin ratio in Western Blots for GAP manipulations**. Data from (A) plx, (B) GAPcenA and (C) Evi5 blots. The data for the GEF Crag is shown in Fig. 2C, where Crag manipulations do affect the level of panRab10. (*G2*: *LRRK2*-*G2019S*; *KD*: kinase dead LRRK2 (*LRRK2*-*G2019S-K1906M*).

