## Supplementary figures and images for "Identification of GEFs and GAPS modulating phosphorylation and abundance of Rab10 by *LRRK2-G2019S* in neurons"

### Supp Fig 1

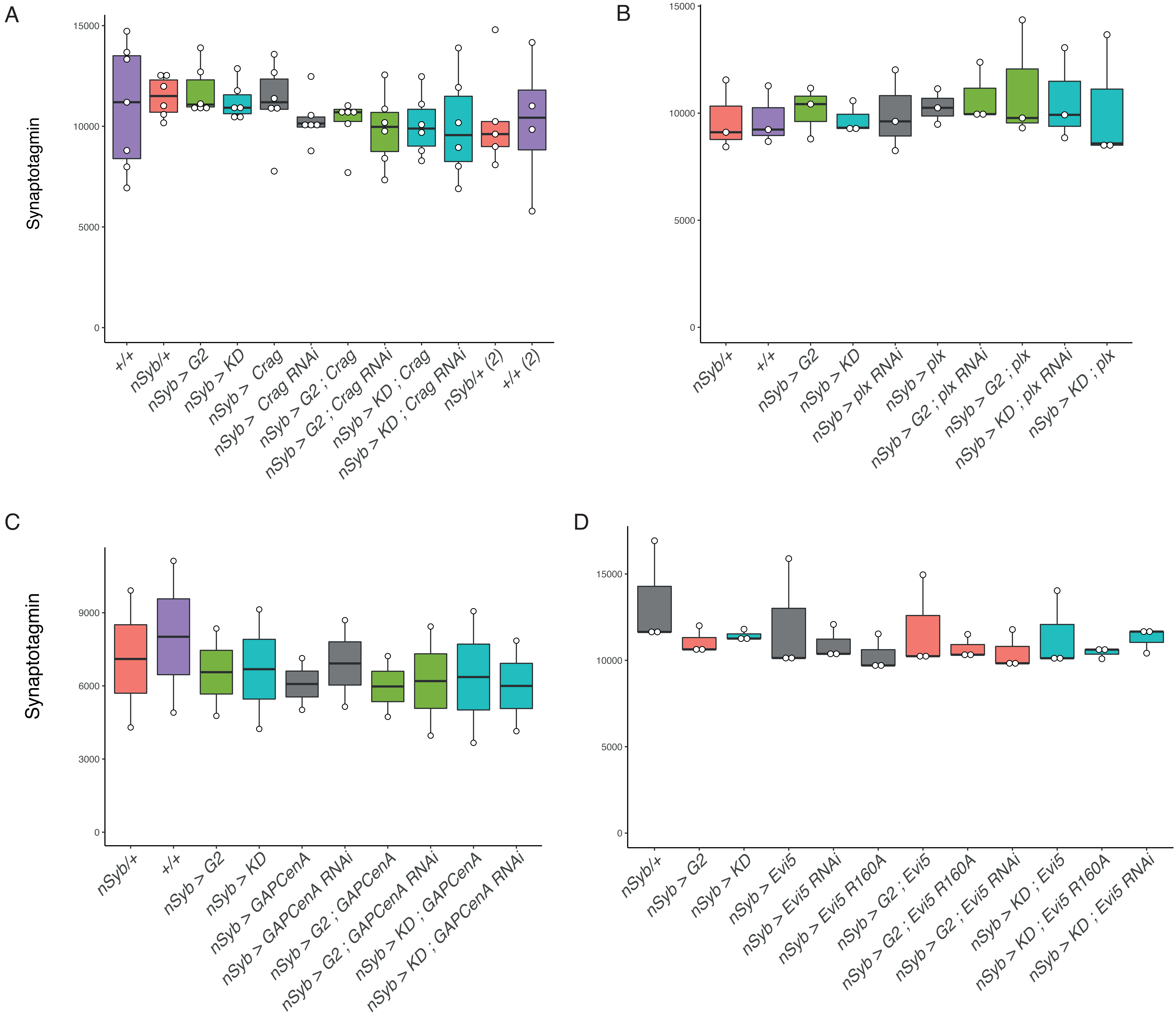

### Supp Fig 2

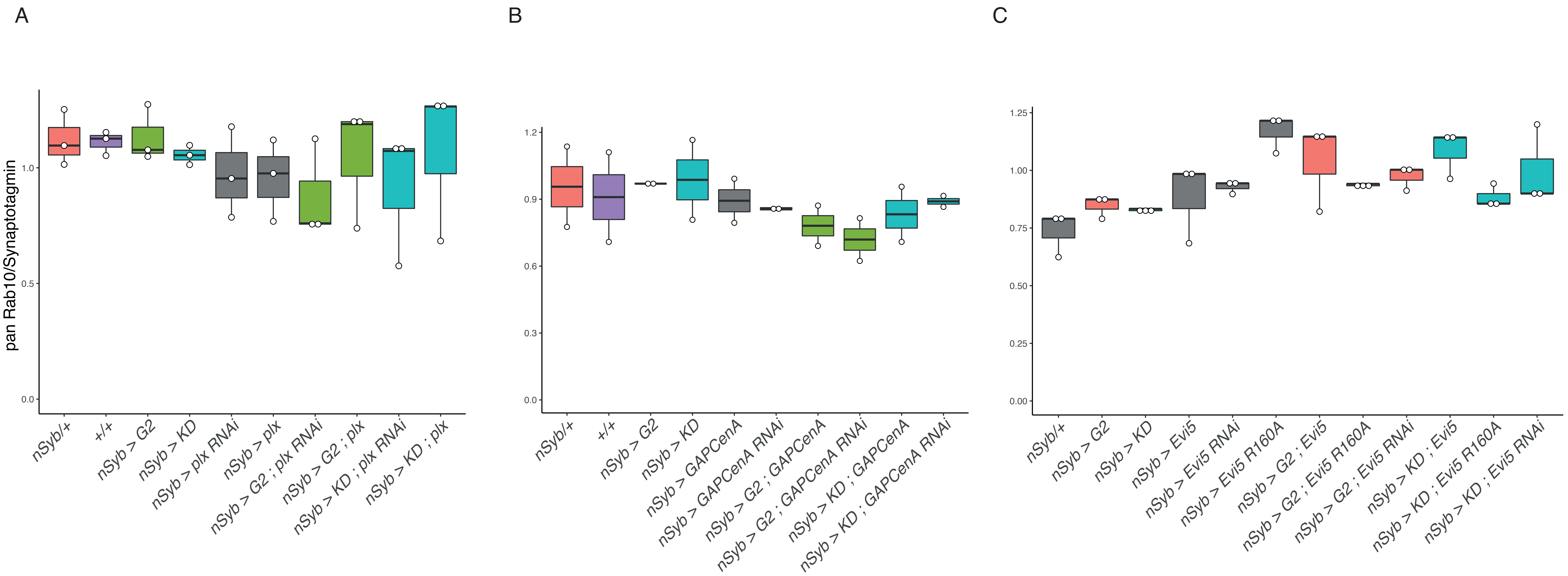
